# Cell size contributes to single-cell proteome variation

**DOI:** 10.1101/2022.10.17.512548

**Authors:** Michael C. Lanz, Lucas Valenzuela, Joshua E. Elias, Jan M. Skotheim

## Abstract

Accurate measurements of the molecular composition of single cells will be necessary for understanding the relationship between gene expression and function in diverse cell types. One of the most important phenotypes that differs between cells is their size, which was recently shown to be an important determinant of proteome composition in populations of similarly sized cells. We therefore sought to test if the effects of cell size on protein concentrations were also evident in single cell proteomics data. Using the relative concentrations of a set of reference proteins to estimate a cell’s DNA-to-cell volume ratio, we found that differences in cell size explain a significant amount of cell-to-cell variance in two published single cell proteome datasets.

## Introduction

Individual cells are the basis of life. It is therefore important to develop techniques that accurately quantify the molecular composition of single cells. Extensive progress examining mRNA composition has been achieved at single cell resolution, helping to catalog diverse cell types in multicellular organisms (1-3). Yet, mRNA sequencing gives an incomplete measurement of the state of the cell because diverse post-transcriptional mechanisms also impact gene expression. For example, the correlation between mRNA and protein amounts is complicated by differing translation and degradation rates (4). Moreover, transcriptomic methods are blind to the diverse set of protein modifications that are often key to activity and function. To address the limitations inherent to measuring only mRNA transcripts, single cell proteomic methods have emerged.

Advances in single cell proteomics are driven by increases in measurement sensitivity from a new generation of mass spectrometers (5). In addition to this increased sensitivity, multiplexed peptide labeling approaches enable the measurement of hundreds and sometimes thousands of proteins from single mammalian cells (6-9). Initial experiments have revealed that the proteomes of single cells are influenced by cell cycle phase (5, 10), though it is unclear which other physiological features underlie cell-to-cell proteome heterogeneity. It is important to measure these and other quantifiable sources of proteome variation to better characterize features that are specific to particular cell types and states.

We recently showed that cell size (*i*.*e*., the DNA-to-cell volume ratio) is an important determinant of proteome content (11). Contrary to the assumption that most cellular components would remain at constant concentration in cells of different sizes, we found widespread, size-dependent changes in the concentrations of individual proteins (**Figure 1A**). These changes in protein concentration likely reflect, to a large extent, the size-dependent changes in the cellular growth rate (12-15). Importantly, a recent proteome analysis of the NCI60 cancer lines revealed a similar pattern of size-dependent changes to the proteome (16). Thus, regardless of cell type, cell size has an important influence on proteome composition and therefore should contribute to the cell-to-cell heterogeneity in the proteomes of single cells.

**Figure 1:**
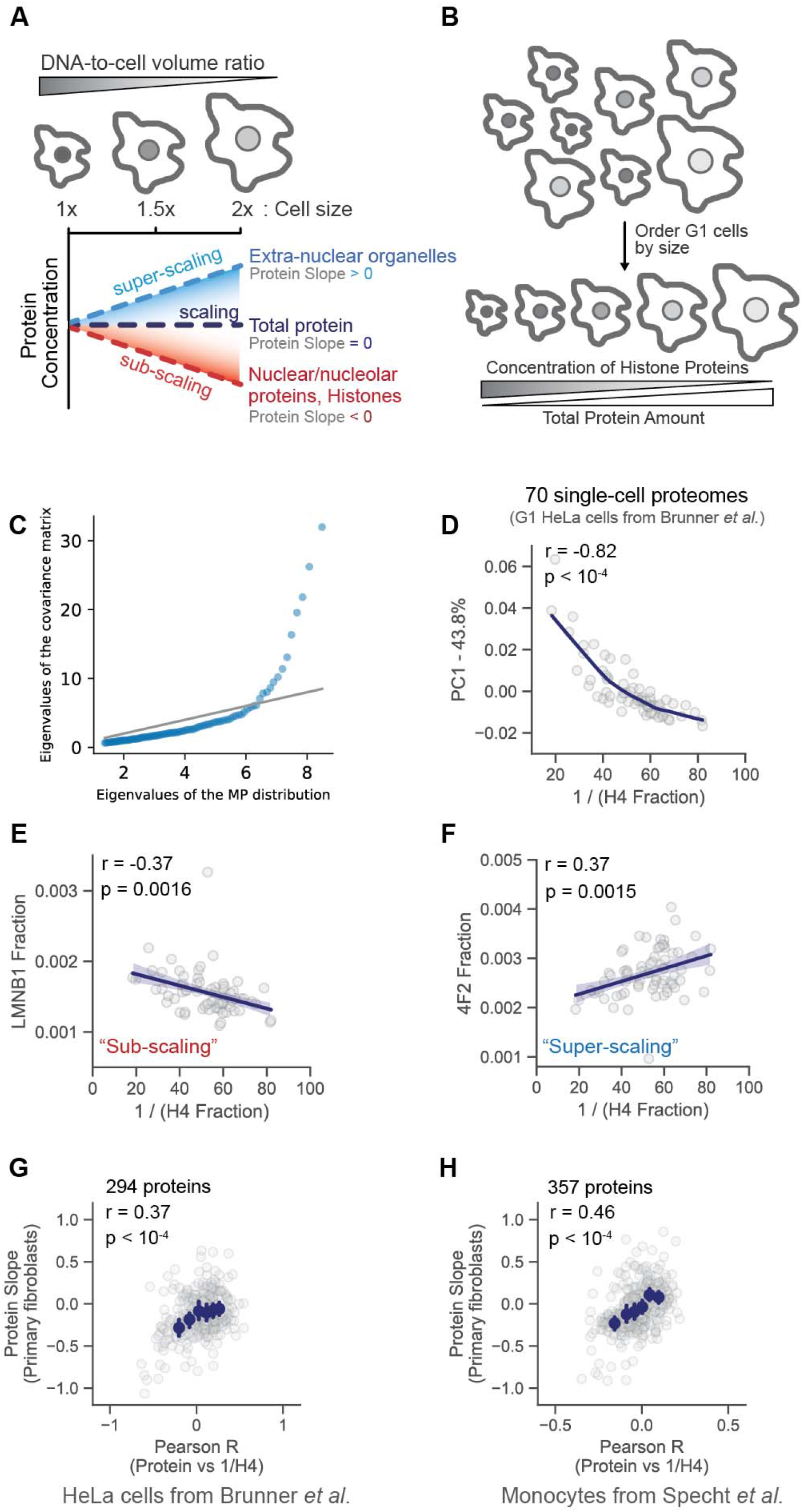
Cell size contributes to variation in the proteomes of single cells. **A)** Proteomes vary with cell size. For example, the amount of histone proteins is maintained in proportion to the genome so that histone concentrations are inversely proportional to cell size. The Protein Slope describes how the concentration of an individual protein scales with cell size (Lanz *et al*., 2022). Proteins with a slope of 0 maintain a constant cellular concentration regardless of cell volume (“scaling”). A slope value of 1 corresponds to an increase in concentration that is proportional to the increase in volume (“super-scaling”), and a slope of -1 corresponds to dilution (concentration ∼ 1/volume; “sub-scaling”). **B)** Schematic illustrating how relative histone protein concentrations can be used as a proxy for cell size in single cell proteomics datasets in which cell size was not measured. **C)** Quantile-quantile plot between the distribution of eigenvalues of the empirical covariance matrix and a sample of the Marcenko-Pastur distribution, which is the distribution expected from uncorrelated, normally distributed random variables. Eigenvalues above the grey identity line indicate the presence of an underlying signal. **D)** PCA analysis of 70 single cell proteomes. Each dot represents the proteome of a G1 cell from Brunner *et al*.. The first principal component is plotted against a proxy for G1 cell size (1/H4 Fraction). The fraction of the proteome represented by histone H4 is H4 intensity / summed intensity of all other proteins. **E)** and **F)** correlation between increasing G1 cell size (1/H4 Fraction) and the relative concentration (*i*.*e*., protein intensity / summed intensity of all other proteins) of two proteins previously found to (E) sub-scale and (F) super-scale with cell size (Lanz *et al*., 2022). **G)** and **H)** A Pearson correlation coefficient was calculated by regressing the relative concentration of each individual protein against a proxy for each cell’s size (1 / H4 concentration), as exemplified in (E) and (F). The r value for each protein from the (G) Brunner *et al*. and (H) Specht *et al*. datasets are plotted against the previously measured Protein Slope value (11). Histone H4 was excluded from the plot. Error bars represent the 99% confidence interval. The plot in (H) was filtered to display the most abundant proteins. Figure S3B depicts an unfiltered version of this analysis.

## Methods

### Data curation

For Brunner *et al*., protein intensities for the individual G1 cells were obtained from PRIDE (ID: PXD024043). G1-labeled columns were extracted from the file named: “20210919_DIANN_SingleCellOutput.pg_matrix.tsv” (DIANN1.8 cell cycle folder). G1 cells without Histone H4 intensity were excluded from the analysis. Also, G1 cells with the fewest number of protein identifications were excluded until a shared set of ∼300 proteins were detected in each single cell. This resulted in the reanalysis of 70 of the 93 G1 cell proteomes (Table S1). For Specht *et al*., a dataframe containing relative protein concentrations for each single cell was downloaded from https://slavovlab.net (“Proteins-processed.csv”). Mock-treated monocytes were extracted from the “Proteins-processed.csv” dataframe using the “sdrf_scope2.tsv” table (Table S2).

### Estimation of cell size

For Brunner *et al*., we estimated the relative cell size for each of the single G1 cells using the “Histone H4 fraction”. To calculate the Histone H4 fraction, we divided the intensity value for Histone H4 (H4_HUMAN) by the summed intensity for all other proteins. To calculate this summed value, we only considered the ion intensity from proteins that were identified in all cells considered for our analysis. We chose a single histone protein, rather than the average of all histone protein, to minimize missing values and therefor maximize the number of cells considered for our analysis. Histone H4 was chosen because a single H4 variant was detected in most cells. The use of other core histone proteins or an averaged value produced similar results (Figure S1). For Specht *et al*., relative cell size was estimated using the relative concentration of Histone H4 (log_2_).

The approach with a single reference protein can be extended to several reference proteins. Namely, we select a small number of reference proteins *n*_*r*_ known to scale significantly with cell size, *i*.*e*., the absolute value of the measured slope *s*_*p*_ for those selected proteins is relatively large. We therefore construct a reference dataset of protein fractions 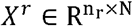 and its associated measured slope values 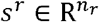 and solve the following regression problem:

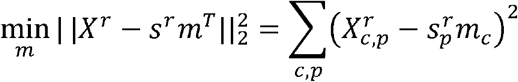

where m ∈ R ^*N*^ is the vector of cell sizes we want to estimate, and the subscripts p, c refer to a specific protein or cell, respectively. In other words, the approach aims to find the cell sizes *m* that, for the reference proteins selected, replicate as closely as possible the previously measured slopes (11). The estimated cell sizes allow the estimation of slopes for all the proteins using standard linear regression.

### Estimation of proportion of variance attributable to cell size

Subtracting the contribution of cell sizes for each protein in the dataset yields a second dataset 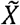 whose total variance is expected to be lower. Indeed, if cell size is a contributor to cell-to-cell proteome variation, removing its effect should decrease the total amount of variance. Denoting the sample covariance matrices of the original dataset and the new dataset with the effect of cell size removed by Σ and 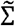, the total variances 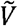 and V-are equal to the sum of the eigenvalues of their respective covariance matrices. Therefore, the amount of leftover variance after removing the effect of slopes is given by 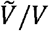. If the contribution of cell size is meaningful, we expect this ratio to be smaller than 1. We can compare this ratio to the leftover variance after removing the first principal component in the dataset, *i*.*e*., *λ*_*max*_ /*V*, where *λ*_*max*_ is the maximum eigenvalue of the sample covariance matrix. The first principal component is the direction in feature space that accounts for maximal variance. Comparing these two metrics reveals the amount of variance contributed by cell size.

### Principal component analysis

A dataframe was created that contained individual proteins as rows with columns corresponding to single G1 cell proteomes. PCA analysis was performed in Python using the sklearn package. Results of the PCA analysis were visualized with Seaborn’s scatterplot.

### Pearson r correlation analysis

For Brunner *et al*., a dataframe containing intensity values of 295 proteins (row) for 70 single cells (column) was converted to intensity fractions. For each cell, the intensity of each protein was divided by the summed intensity of all proteins to calculate each protein’s proteome fraction, an estimate of a protein’s relative concentration. A Pearson r correlation (python’s scipy package) was calculated by regressing the relative concentration of each individual protein against a proxy for each cell’s size (exemplified in Figure 1E and 1F). For Specht *et al*., we used the log_2_ ratio values published by the authors, so the r value was derived from a regression between the relative protein concentration (log_2_) versus the relative Histone H4 concentration (log_2_). Only the most abundant ∼350 proteins were considered for Figure 1H (filtered by peptide detections in our own dataset). Our analysis of the entire Specht *et al*. dataset can be found in Figure S3 and Table S2.

## Results

To investigate whether cell size can explain cell-to-cell variations in proteome content, we reanalyzed data from two recently published single-cell proteome datasets (5, 6) (**Table S1 and S2**). One of these utilized Bruker’s ultra-high-sensitivity timsTOF SCP (label-free DIA) to measure the proteomes of single HeLa cells that were proceeding through the cell cycle after being synchronized (5). The authors distinguished the cell cycle phase of single cells based on their measurements. To disentangle cell size and cell cycle-related effects, we only considered the proteomes of G1-enriched single cells for our analysis. An eigenvalue analysis of this set of G1 cell proteomes found that the top eigenvalues of the covariance matrix deviate significantly from the Marcenko-Pastur distribution (17), *i*.*e*., the distribution of eigenvalues for datasets with no latent variables (**Figure 1C**). This means there is a significant signal in our data despite the noisy nature of single cell proteomics data. To crudely approximate the relative size of each cell, we used the histone proteins because their amount is proportional to the amount of DNA (11, 18-20). Smaller cells therefore possess proportionally higher concentrations of histone proteins than larger cells (**Figure 1B**). We used the fraction of total ion intensity represented by Histone H4 as representative of the inverse of the cell volume (a proxy for cell size). We performed principal component analysis (PCA) on 70 G1-enriched single-cell proteomes from Brunner *et al*., reasoning that proteins with cell size-dependent abundances could help explain the variance in these cells. The fraction of total ion signal attributable to histone H4 significantly correlated with the first principal component indicating the importance of cell size in contributing to cell-to-cell proteome variation (**Figure 1D**). Other core histone proteins produced similar results (**Figure S1**). In contrast, substituting a histone protein for a common housekeeping enzyme, PGK1, whose concentration is expected to be independent of cell size (11), did not produce a significant correlation (**Figure S2**).

To further explore the relationship between single cell proteome variation and cell size, we calculated Pearson coefficients (r) for each protein from the correlation between its relative protein concentration and Histone H4, a proxy for cell size (**Figure 1E and 1F**). We then correlated these r values with the protein concentration size-dependence previously reported by Lanz *et al*. (**Figure 1G**). Concentration size-dependence was calculated as a protein slope. In brief, the protein slope is calculated from a linear regression between the log_2_ of an individual protein’s concentration and the log_2_ of the cell volume. Thus, a protein slope value of 0 describes proteins for which concentration does not change with cell volume (scaling), a protein slope value of 1 describes proteins for which concentrations increase proportionally to cell volume (super-scaling), and a protein slope value of -1 describes proteins that are perfectly diluted by cell growth such that their concentration is inversely proportional to cell volume (sub-scaling). The Pearson r value correlating concentration and histone H4 derived from single cells was correlated with the previously published protein slope values (**Figure 1G)**. Having established that cell size influences variation in one single cell proteomics dataset, we next sought to examine the robustness of this result in a second dataset. To do this, we repeated this analysis on a second dataset generated using a different single-cell proteomic platform (6). Using SCoPE2, Specht *et al*. distinguished individual monocytes that were or were not differentiated into macrophages. Like the HeLa cells measured by Brunner *et al*. (**Figure 1G**), Pearson regression analysis of single monocyte proteomes produced r values which significantly correlated with the protein slope values (**Figure 1H)**. Taken together, these correlations strongly support the hypothesis that variations in cell size measurably contribute to single-cell proteome variation.

Since single cell proteomics measurements are noisy, we anticipate that there is significant noise in our estimate of cell size using only histone H4. We therefore sought to derive a more robust method for measuring the size-dependent proteome variation in single cell proteome datasets. To do this, we decided to use more than one reference protein to approximate a cell’s size. We selected a subset of reference proteins and reconstructed cell sizes based on a least-squared-error heuristic (**Figure 2A**). Under this framework, the estimated cell size distribution is the one that most closely reproduces the set of measured protein slopes for this subset of proteins. Assuming that protein-to-protein noise is uncorrelated, using more than one reference protein ensures the impact of measurement noise is reduced. The reference proteins were chosen based on their (i) large absolute measured slope value and (ii) large median correlation coefficients with other reference proteins. These criteria ensure that the reference proteins encode meaningful variations from which a signal can be extracted.

**Figure 2:**
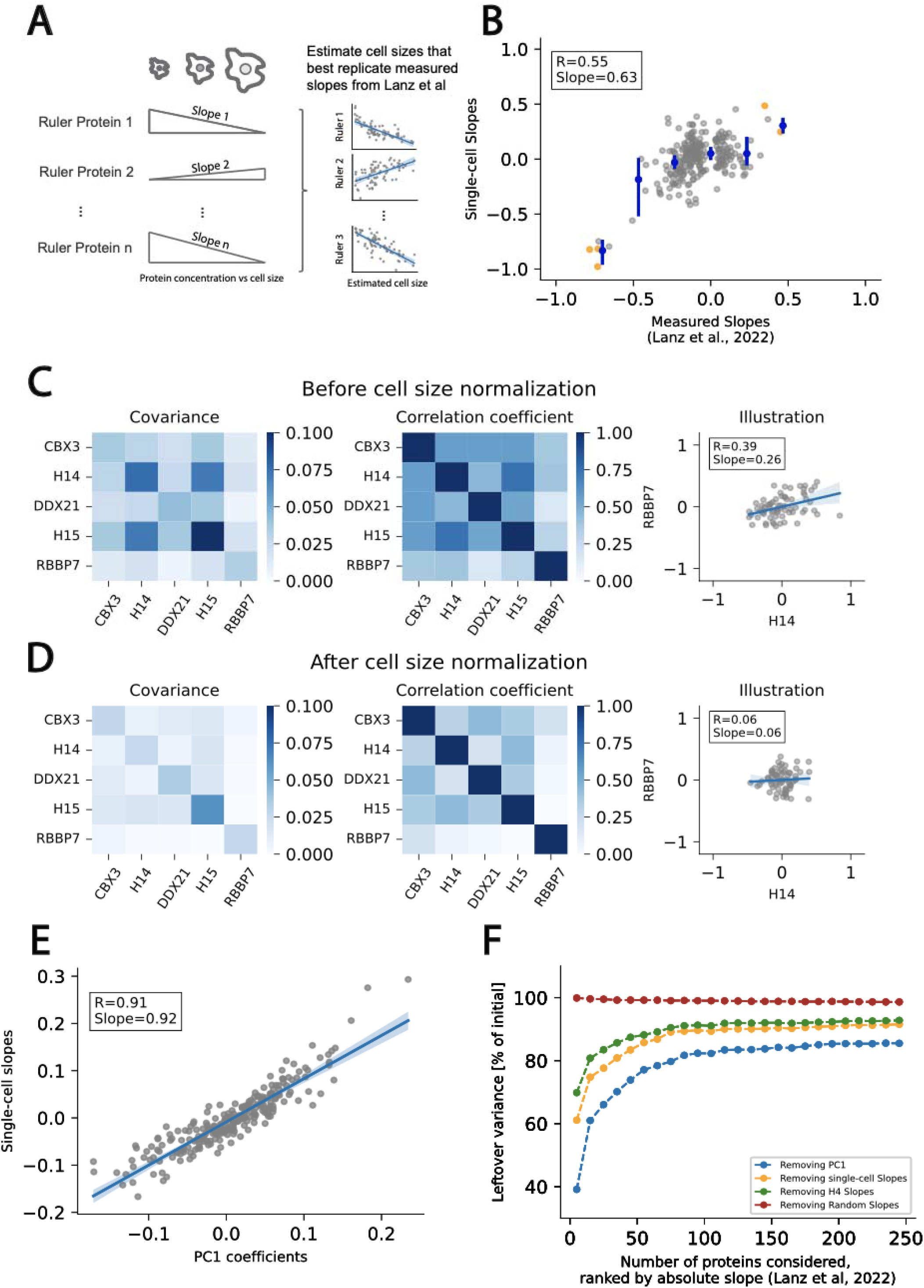
Robust estimation of cell size used to determine the size-dependent variance in single-cell proteomics data. **A)**Schematic illustrating the methodology to estimate cell size. We first select a small subset of reference proteins, like histone H4, whose concentrations were shown to be strongly size-dependent (Lanz et al., 2022). Using these reference proteins, and their corresponding size slope values, we performed a least-squares regression to estimate the size of each cell. **B)**Having estimated the size of each cell, we then calculate the size slope for each protein in our single cell proteomics data sets. Comparison of slopes estimated via the approach described in this paper and those measured previously (11). Orange dots denote reference proteins and blue dots with error bars denote binned values. **C and D)** Comparison of protein concentration covariance and correlation in the initial dataset (**C**) and after removing the estimated effect of cell size (**D**) for a set of 5 proteins with large absolute measured slopes. Removing the estimated effect of cell size reduces the covariance and the correlation coefficient between protein pairs. We illustrate this effect with a given protein pair. **E)** Relationship between estimated slopes and the coefficients of the first principal component. Both quantities are very close to each other, indicating the estimated slopes approximate the direction of maximum variance in the dataset. **F)** Amount of variance leftover after removing the first principal component (blue), the estimated effect of cell size from measured slopes (orange), the effect of H4 only (green), and the estimated effect of cell size from random slopes (red). The number of proteins included in the analysis (x-axis) was gradually increased based on protein absolute slopes. For example, if 50 proteins were included in the analysis, this set contains the 50 proteins with the highest absolute slopes. The maximum amount of variance removed by cell size is bounded that removed by the first principle component (PC1 blue).

Based on our single reference protein analysis in **Figure** 1, the scaling behavior of proteins is expected to be qualitatively conserved across cell types. Using the reconstructed cell size distribution and the single-cell proteome measurements, we compute *single-cell slopes* for each of the remaining proteins in the dataset, which were well correlated with the previously measured slopes (**Figure 2B**). These results are qualitatively similar to the analysis shown in **Figure 1**, but quantitatively more robust due to the use of five reference proteins to estimate cell size. Indeed, a smaller number of rulers yields larger variations in the estimated single cell slopes and cell size distributions (**Figure S4**). The correlation between estimated and measured slopes does not vary significantly when more than five rulers are used (**Figure S5**).

Having found that cell size contributes to single cell proteome variation, we next sought to quantify the extent of this contribution to the total variation in single cell proteomics data. To explore this contribution, we first subtracted the estimated contribution of cell size to each protein concentration from our data. This resulted in a significant decrease in the covariance and correlation coefficient among a set of proteins whose concentrations varied with cell size (**Figure 2C-D**). To assess whether the estimated cell size distribution is the primary source of variance in the dataset, we compared the first principal component coefficients with the imputed *single-cell* slopes (**Figure 2E**). The strong correlation demonstrates that the estimated slopes closely align with the direction of maximum variance. Consistent with this observation, no correlation is observed with the second PC coefficients (**Figure S6**). Finally, the imputed cell size distribution allowed us to estimate the proportion of variance in the dataset that is attributable to cell size (see Methods; **Figure 2F**). The variance due to cell size differences is comparable to the variance attributed to the first principal component, which sets the upper bound for removable variance by a single linear transformation. While using histone H4 as a single reference protein does remove some variance in the dataset, using more reference proteins increases the amount of variance that can be accounted for. We note that using protein slopes derived from the measurement of another cell type (11) yields generally similar results (**Figure S7**). In contrast, using a set of randomly generated protein slopes does not remove any variance in the dataset.

## Discussion

While our data demonstrate that cell size is a major contributor to variation in single cell proteomes extracted from cells of the same type, it is important to note that relationship may be more complex when datasets contain very different types of cells, as was recently demonstrated in a melanoma cell line (10). Nevertheless, even in these complex assemblies of cells, we still anticipate there to be some size-dependent signal because several different cell lines exhibit similar size scaling across their proteomes (11, 16). We also note that differences in ploidy across cell types may have a big effect on the proteome. For example, a comparison of the proteomes of 2N, 4N, and 8N cells revealed that they were highly similar despite the near 4-fold increase in cell size (11). Thus, what we refer to as size scaling is more accurately attributed to changes in the cell volume-to-DNA ratio.

In summary, we re-analyzed the proteome heterogeneity in single cell datasets reported by two independent groups using different single-cell preparation and measurement platforms. In both cases, we found that differences in cell size substantially contribute to variance in single cell proteomes. Remarkably, the effects of cell size trended in agreement with a recent report of cell size-dependent changes to the proteome that were measured in bulk for different types of cells (11). Taken together, these analyses support the conclusion that differences in cell size will account for a significant amount of proteome heterogeneity in single cells. We therefore recommend accounting for differences in cell size in future analyses of single cell proteomes.

## Acknowledgements

We thank Martin WÜhr and members of the Skotheim and Elias labs for feedback on the manuscript. We also thank the authors of Brunner *et al*. and Specht *et al*. for collecting and curating their single cell data in a manner that facilitated our reanalysis.

Funding: Chan Zuckerberg Biohub (collaborative postdoctoral fellowship M.C.L.; investigator award J.M.S.); NIH P01 CA254867.

## Data Availability

This study analyzed previously published data.

## Author contributions

M.C.L. and L.V. performed data analysis and wrote the manuscript with J.E.E. and J.M.S, who supervised the study.

## Supplementary information

**Figure S1:**
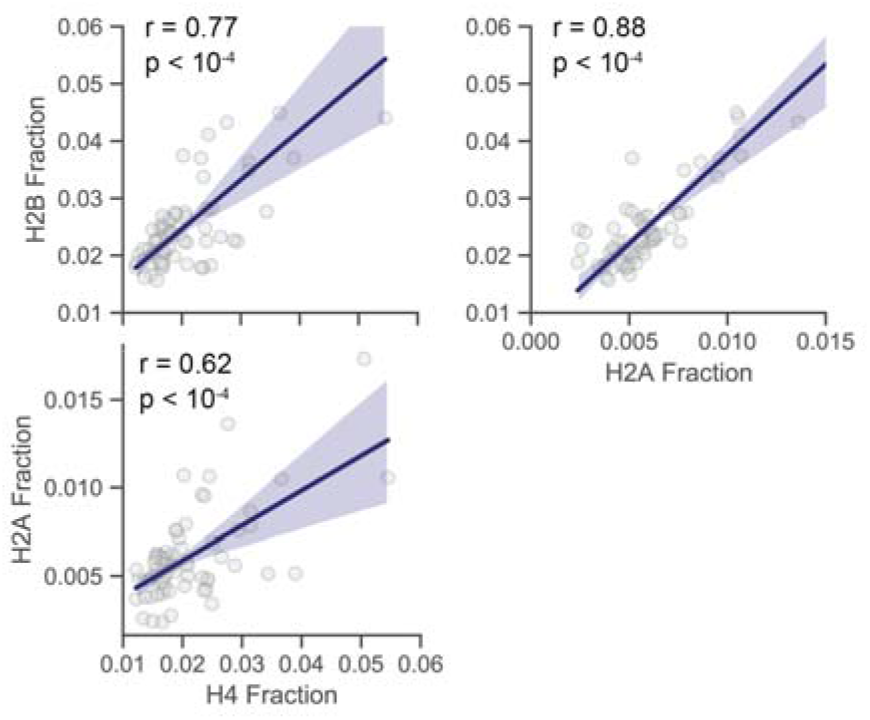
Multiple core histones can be used to estimate cell size. The concentration of Histone H4 correlated with the concentration of other core histones in single cells. A regression line is plotted in dark blue with 95% confidence intervals. Pearson r value and its associated p-value are shown.

**Figure S2:**
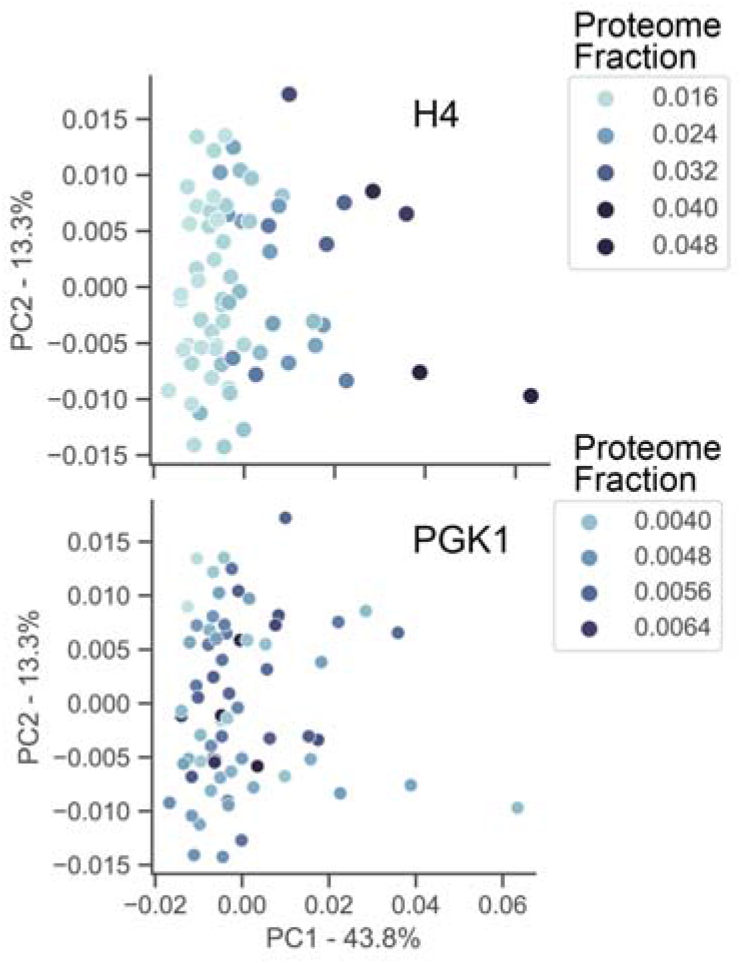
Correlation of principal components with the relative concentration of histone H4 and PGK1, a protein whose concentration is not expected to change with cell size. Plot of the first two principal components from a PCA analysis of 70 single cell proteomes. Each dot represents the proteome of a G1 cell from Brunner *et al*. and its color indicates the fraction of the proteome represented by either H4 (top) or PGK1 (bottom).

**Figure S3:**
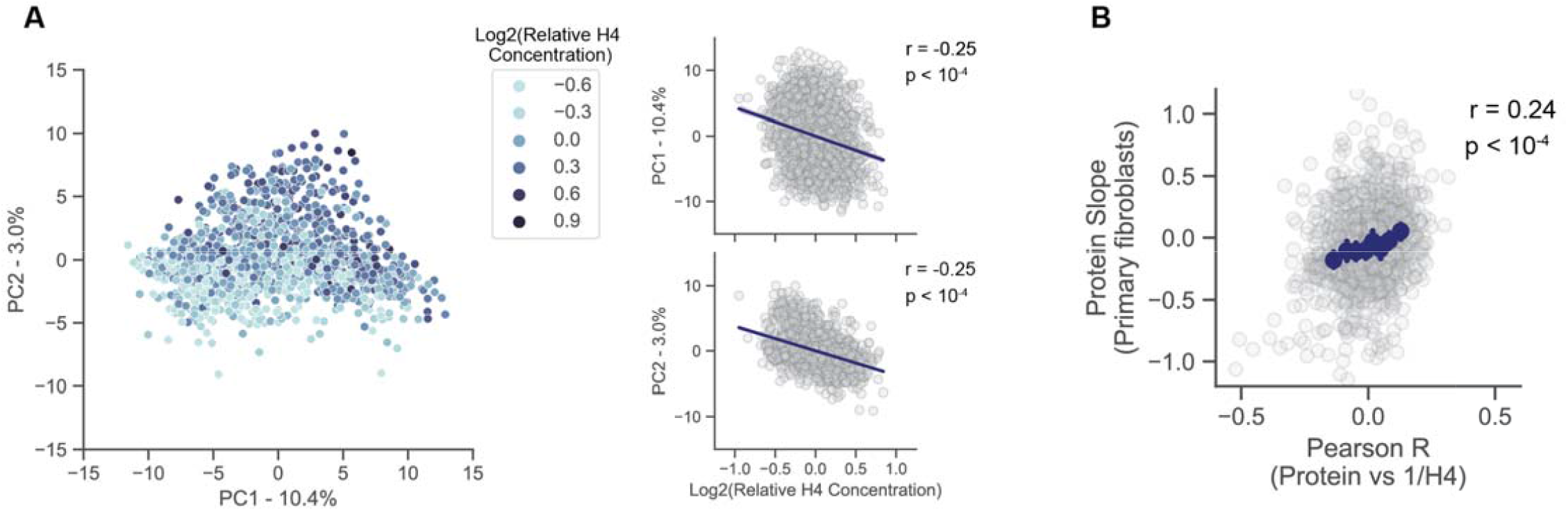
Reanalysis of single cell proteomics data from Specht *et al*. **A)** PCA analysis of 1490 single cell proteomes from Specht *et al*. Proteins were quantified using tandem mass tags, so cell size was estimated using the relative concentration of MS2-level reporter ions for Histone H4. Each dot represents a proteome and its color indicates the relative H4 concentration. Correlation between the relative histone concentrations and PC1 and PC2. A regression line is plotted in dark blue with 95% confidence intervals. Pearson r value and its associated p-value are shown. **B)** A Pearson correlation coefficient was calculated by regressing the relative concentration of each individual protein against a proxy for each cell’s size (histone H4 concentration). The r value for each protein in the Specht *et al*. dataset is plotted against the Protein Slope value (Lanz *et al*., 2022). Histone H4 was excluded from the plot. Error bars for the binned data represent the 99% confidence interval of the mean. In Figu e 1H, only the most abundant proteins are depicted. All proteins identified by Specht *et al*. are included.

**Figure S4:**
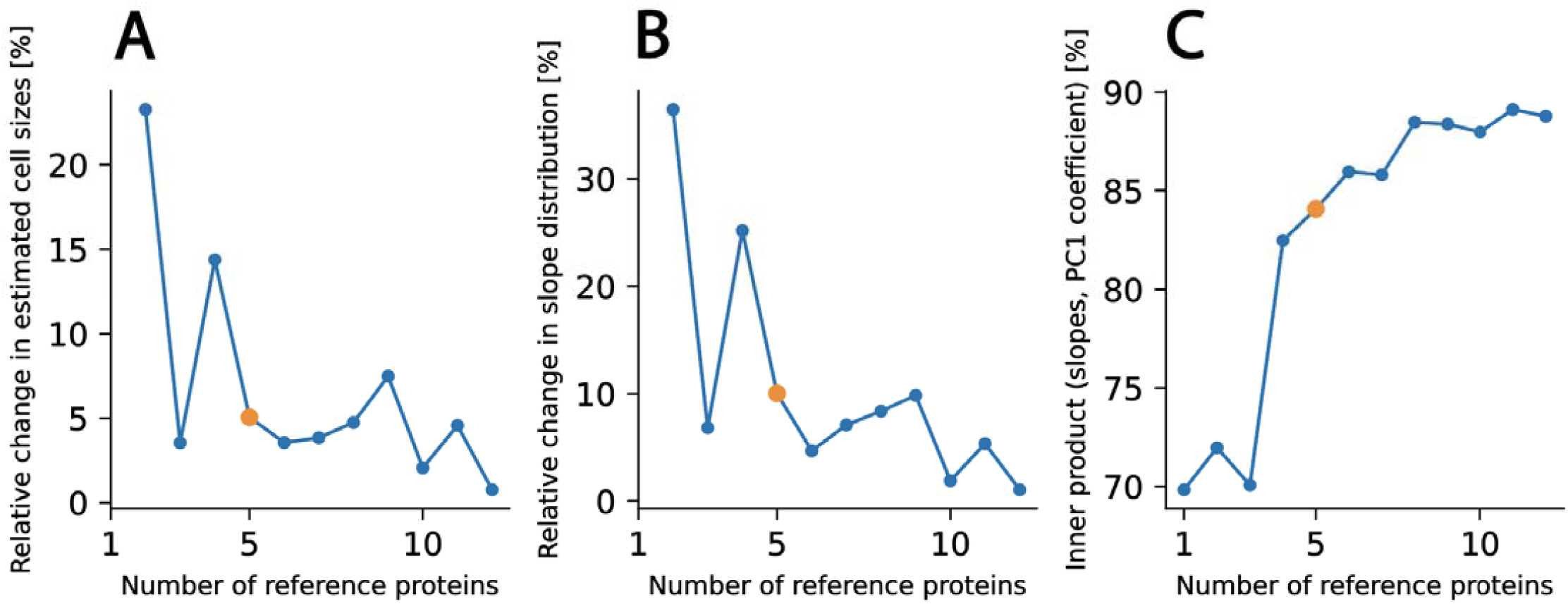
Metrics used to determine the number of reference proteins. The orange dot represents the number of reference proteins that is presented in the main text. Relative change denotes the value for n reference proteins minus the value for n-1 reference proteins divided by value for n reference proteins. **A)** Relative change in inferred estimated cell size distribution as more reference proteins are added. **B)** Relative change in estimated slope distribution as more reference proteins are added. **C)** Normalized inner product between the slopes and the PC1 coefficient as a function of the number of reference proteins. Reference proteins were discarded from the dataset before computing this metric. We chose 5 reference proteins because beyond this number changes produced only marginal differences.

**Figure S5:**
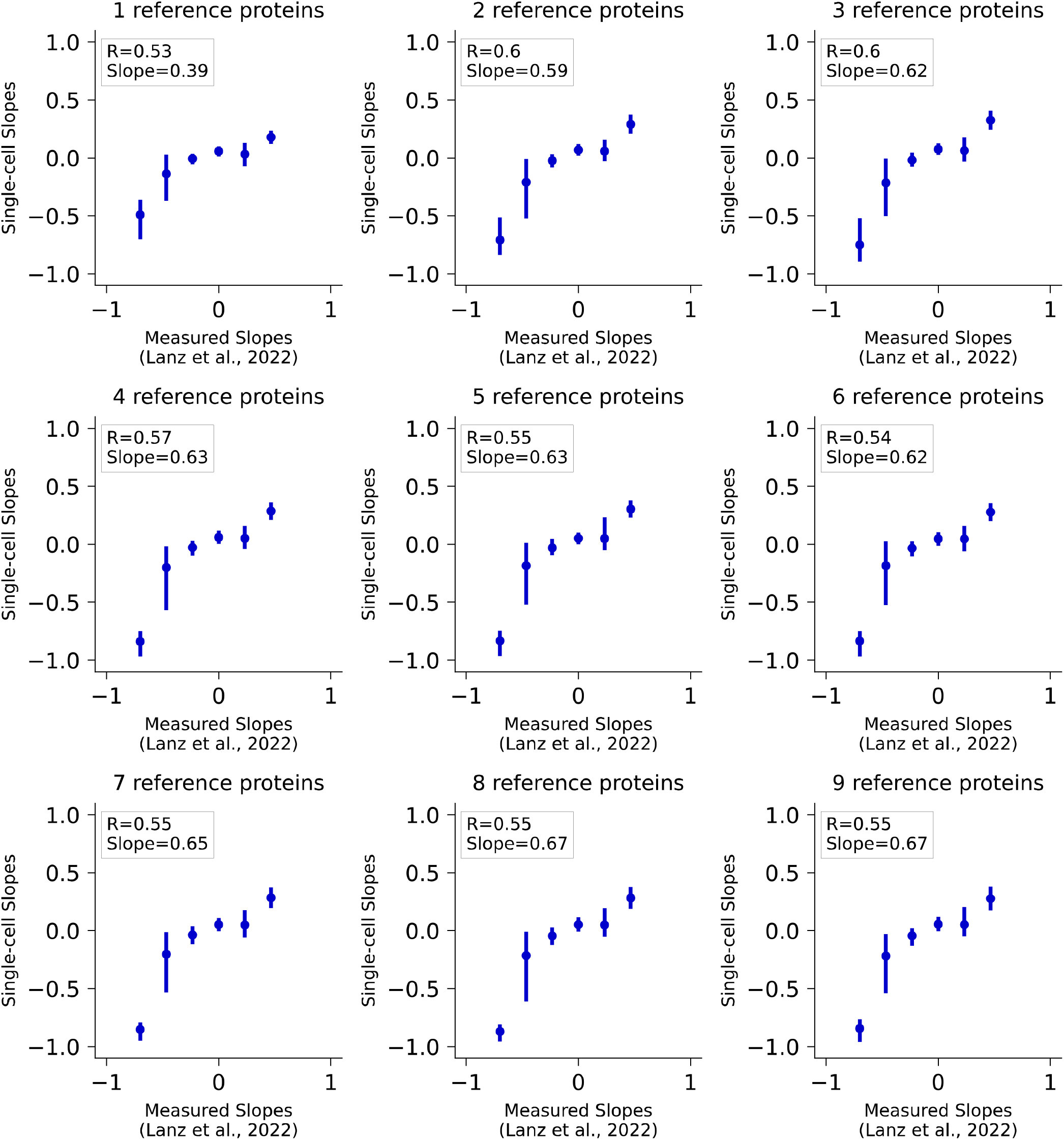
Impact of increasing number of reference proteins on single cell slopes. Each panel reports the slopes estimated with a given number of reference proteins. Orange dots denote the proteins used as reference and blue dots denote binned data and associated confidence intervals.

**Figure S6:**
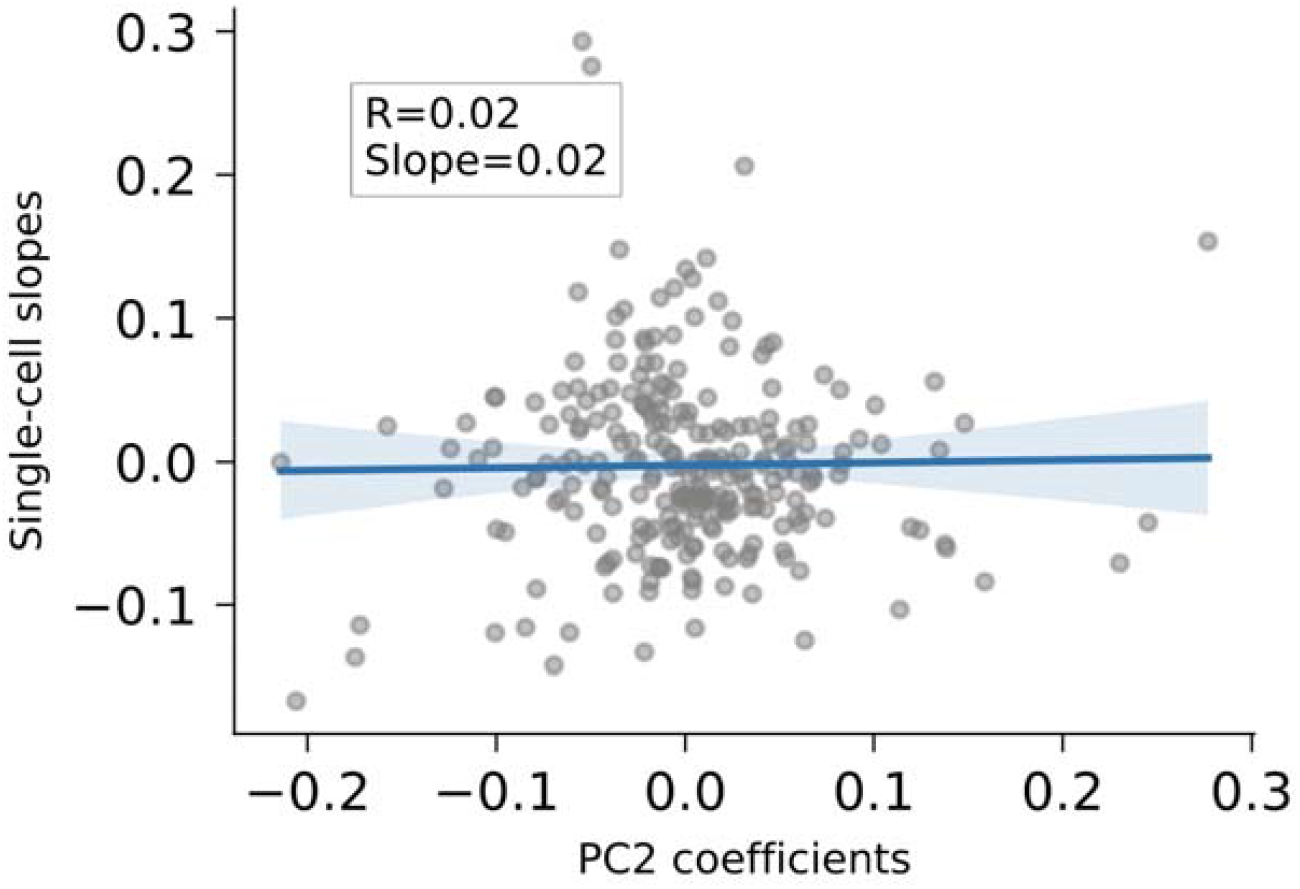
Correlation of estimated single-cell protein slopes with the coefficients from PC2 from a PCA analysis of *Brunner et al*. as shown for PC1 in Fig. 2E.

**Figure S7:**
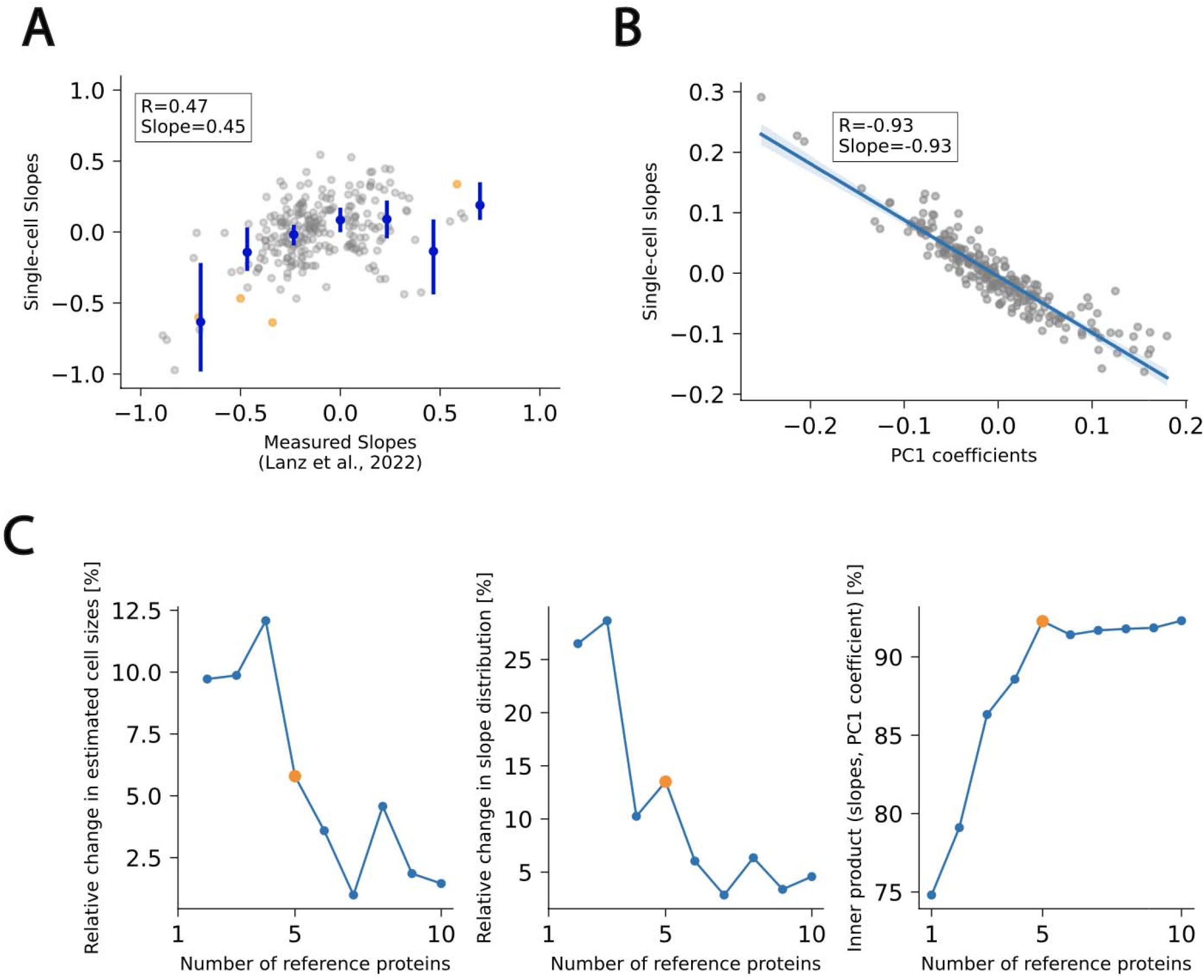
Analysis of the *Brunner et al*. dataset using HLF protein slopes from *Lanz et al*. rath RPE1 protein slopes. **A)** Comparison of protein slopes estimated via the approach described in this paper and those measur *Lanz et al*. Orange dots denote the proteins used as reference and blue dots denote binned data and associated confidence intervals. **B)** Relationship between estimated slopes and the coefficients of the first principal component. **C)** Relevant metrics with increasing number of reference proteins as in **Figure S4**.

